# Extracellular Nanobody Screening using Conformationally Stable GPCR

**DOI:** 10.1101/2025.03.25.645380

**Authors:** Xin Zhang, Kaixuan Gao, Jia Nie, Hengyu Meng, Xiaoou Sun, Jiawei Zhao, Xiangyu Liu

## Abstract

G protein-coupled receptors (GPCRs) are prominent drug targets that have attracted intensive efforts in drug screening. Binding-based screening methods for GPCR ligands often require conformationally stable, purified receptors. However, obtaining large quantities of GPCRs in stable states, particularly with unoccupied extracellular ligand-binding pockets and especially in their active conformations, remains challenging due to the inherent dynamic nature of these receptors. To address this challenge, we propose a universal approach for stabilizing GPCRs in specific conformations. Using the M1 muscarinic acetylcholine receptor (M1R) as a model, we successfully stabilized M1R in its active conformation through de novo design of a fusion protein. We screened a synthetic yeast display library of nanobodies against both the stabilized active-state and previously reported inactive-state M1R, identifying several nanobodies that specifically recognize each conformation. This method not only facilitates the stabilization of GPCRs in desired states but also provides valuable tools for developing more selective therapeutic agents, enhancing drug discovery efficiency and specificity.

## Introduction

GPCRs are central mediators of signal transduction across cell membranes, regulating nearly every aspect of human physiology. Their pivotal role in modulating physiological processes has made GPCRs prime therapeutic targets. Notably, approximately 34% of all FDA approved drugs function by modulating GPCR activities (1). Despite extensive pharmacological and structural studies that have elucidated the mechanisms by which ligands stimulate or inhibit these receptors, drug discovery efforts targeting GPCRs still face significant challenges. A particularly pressing challenge is the development of conformation-selective and subtype-selective ligands (2, 3).

The conformational flexibilities of GPCRs complicates the direct enrichment of conformation specific ligands. GPCRs typically exist in a spectrum of conformations between two primary states: the inactive state and the active state (4). The prominent conformational change occurs in the intracellular domain, particularly in Transmembrane Helix 6 (TM6), while the dynamics in the extracellular domain, which contains the orthosteric pocket, are relatively minor (5, 6). These subtle conformational changes do not prevent the binding of antagonists or agonists but instead modulate their affinity for receptors in different states (7). Identifying ligands by recognizing these subtle conformational changes is crucial for discovering function-specific ligands.

Binding-based experimental screening techniques that utilize purified receptors, such as DNA-encoded libraries (DEL), yeast surface displayed libraries, and phage displayed libraries, present a promising approach encompassing chemical space exploration and conformational selection (8–11). A pivotal challenge in this context is obtaining conformation-stable purified receptors, particularly in their active states, which are thermodynamically unstable. Molecular dynamics studies on receptors such as the beta-2 adrenergic receptor (β2AR), µ-opioid receptor (MOR), and muscarinic acetylcholine receptor 2 (M2R) have demonstrated the instability of active conformations (12–16). Single-molecule studies reveal that under apo conditions or when bound with antagonists or partial agonists, these receptors predominantly remain in the inactive state. Even in the presence of full agonists, the receptor fluctuates between active and inactive states (17, 18). This transient active state poses a significant obstacle for agonist screening. To address this issue, full agonists, G proteins, or G protein mimics have been employed to stabilize the active state during screening (19–21). Although these strategies achieve some success, they also come with limitations. Full agonists occupy the orthosteric pocket, making it impossible to screen for other orthosteric agonists. Moreover, the resulting receptor-G protein or receptor-G protein mimics complexes are often unstable, with dissociation disrupting the screening process (22). Therefore, there is an urgent need for innovative strategies to design conformation-stable receptors that can effectively overcome these challenges.

Muscarinic acetylcholine receptors represent a prominent class of receptors that have attracted considerable attention as promising targets for addressing complicated neuropsychiatric disorders, including Alzheimer’s disease, schizophrenia, and Parkinson’s disease (23–26). Despite extensive efforts, only one drug based on the cholinergic mechanism, xanomeline combined with peripheral antagonist trospium chloride (KarXT), received FDA approval in 2024 for the treatment of schizophrenia (27, 28). A major hurdle has been the occurrence of cholinergic adverse effects, often resulting from the off-target activation of other muscarinic acetylcholine receptor subtypes, leading to the discontinuation of most clinical trials (25). Consequently, over the past decade, research in both academia and industry has focused on enhancing the subtype selectivity of muscarinic acetylcholine receptor modulators. Antibodies offer a promising solution to the challenges of developing subtype-selective ligands, given their ability to engage broader epitopes across receptor surface, thereby facilitating more selective binding (29–31). However, the development of antibody-based modulators targeting the muscarinic acetylcholine receptor family remains largely unexplored.

Previously, we reported the de novo design of a fusion protein that locks M1R in an inactive state (32). In this study, we extended our design strategy to stabilize M1R in its active state. This approach resulted in a GPCR that remains in the active conformation even in the absence of agonist binding. Leveraging this design, we screened a series of nanobody targeting M1R. Structural and pharmacological analyses revealed that these nanobodies function as active-state selective nanobodies or inactive-state selective nanobodies by interacting with the extracellular pocket of M1R. Together, our findings validate the conceptual feasibility of screening extracellular nanobodies targeting GPCRs using conformation-locked GPCRs and offer new tools for drug discovery and mechanistic studies of M1R.

## Results

### Generation of conformationally stable M1R in active state

With the advancements of generative protein design technology, it has become feasible to create a wide variety of proteins with shapes precisely tailored to specific requirements. A hallmark of GPCR activation is the outward movement of TM6. Our objective is to design de novo fusion proteins that connect TM5 and TM6, locking TM6 into either an inward or outward displacement, which stabilizes the receptor in its inactive or active conformation, respectively (Fig. 1A). Previously, we reported our approach for designing an inactive-state GPCR using a click fusion protein named Clip1, which is rigidly linked to TM5 and TM6 to restrict the dynamic of TM6. For designing an active-state GPCR, the α5 helix of the G protein is employed to lock the outward movement state of TM6. Studies have demonstrated that the α5 helix of the G protein alone can stabilize the active state of the receptors such as β2-adrenergic receptor (β2AR) and rhodopsin without the requirement for the intact G protein or engineered miniG protein (33–35). To implement this, the α5 helix was linked to the C-terminus of the receptor via a 2×GGSGG linker, with a proline residue added at the N-terminus of the α5 helix of G protein to promote helical formation (36). Meanwhile, a de novo designed fusion protein containing a binding pocket for the α5 helix of G protein was fused between TM5 and TM6 to stabilize the interaction between the receptor and the α5 helix of G protein (Fig. 1B). The cryo-EM structure of the active-state M1R bound to the G_11_ protein (PDB: 6OIJ) served as the reference structure model. Initially, the receptor chain and the α5 helix of the G_11_ protein (residues 335-359, ENIRFVFAAVKDTILQLNLKEYNLV, G_11_CT) were retained from the original model. Subsequently, the backbone of the fusion protein was generated between TM5 and TM6 using RFdiffusion, forming a suitable binding pocket to stabilize the position of the G_11_CT. Sequence design was performed using ProteinMPNN and structure prediction was conducted using AlphaFold2. Two sequences with high pLDDT scores were selected and synthesized, named M1R-G_11_CT and M1R-G_11_CTb (Fig. 1C). While the two proteins share the same scaffold, their surface charge distributions exhibit significant differences (SI Appendix, Fig. S1 A and B).

**Fig. 1.**
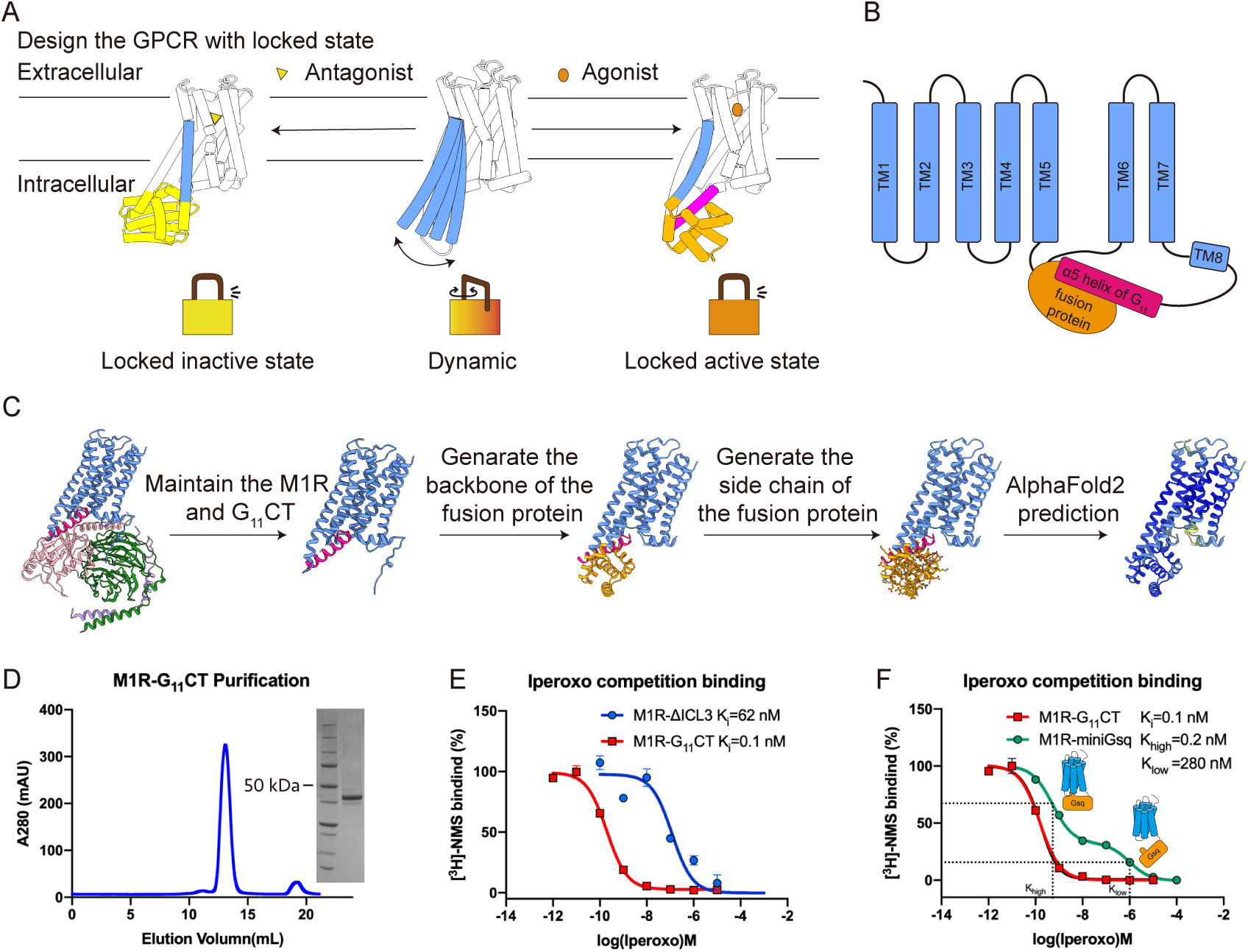
Design and biochemical characterization of M1R-G_11_CT. A) Schematic representation of stabilizing GPCRs in either the inactive state or active state, with TM6 constrained in an inward or outward conformation through de novo designed fusion protein. (B) Schematic representation of the M1R-G_11_CT construct, featuring G_11_CT fused to the C-terminus of receptor and the designed fusion protein inserted between TM5 and TM6 that accommodated the G_11_CT. (C) Overview of the design process for the fusion protein of M1R-G_11_CT. (D) SEC profile and corresponding SDS-PAGE analysis of M1R-G_11_CT. (E) Competition binding assays comparing M1R-G_11_CT and M1R-ΔICL3 with iperoxo demonstrate that the engineered M1R-G_11_CT, which exhibits enhanced agonist affinity, stabilizes the receptor in its active state. (F) Competition binding assays of M1R-G_11_CT and M1R-miniGsq with iperoxo demonstrate the homogeneity and stability of M1R-G_11_CT. Data are represented as mean ± SEM (n=3).

### Biochemical characterization of M1R-G_11_CT

Cell surface expression analysis revealed that M1R-G11CT and M1R-G11CTb demonstrated comparable expression levels to both M1R-Clip1 and M1R-ΔICL3, the latter representing a widely utilized construct featuring a truncated intracellular loop 3 (ICL3) (SI Appendix, Fig. S1C). And the yield of purified M1R-G_11_CT and M1R-G_11_CTb protein reached approximately 500 µg per liter of cell culture. Notably, M1R-G_11_CT and M1R-G_11_CTb were purified in the absence of any ligand, similar to M1R-Clip1 (32). The purified proteins displayed excellent behavior during size-exclusion chromatography (SEC), as evidenced by the sharp and symmetrical peaks with minimal aggregation (Fig. 1D and SI Appendix, Fig. S1D). These characteristics suggest that M1R-G_11_CT and M1R-G_11_CTb proteins are intrinsically well-folded and homogeneous without the need for additional stabilizing ligands.

Previous studies have shown that M1R-ΔICL3 maintains functional properties similar to those of the wild-type M1R, including ligand binding profiles and receptor signaling capabilities (6). Isotopic ligand binding assays were performed to validate the biochemical characterization of these constructs. Saturation binding assays were conducted to determine the affinity of [³H]-N-methylscopolamine (NMS) for these constructs. M1R-G_11_CT exhibited a Kd value of 1.4 nM (SI Appendix, Fig. S1E), comparable to the previously measured 1.1 nM for M1R-ΔICL3, indicating similar affinities for the antagonist NMS (32). Additionally, radioligand competition binding assays were conducted to evaluate the affinities of the agonist iperoxo and the antagonist atropine in these constructs. Compared to M1R-ΔICL3 (Ki=1.9 nM), M1R-G_11_CT showed a similar binding affinity for atropine (Ki=0.6 nM) (SI Appendix, Fig. S1F). Remarkably, M1R-G_11_CT displayed an approximately 600-fold higher binding affinity for the agonist iperoxo relative to M1R-ΔICL3 (Fig. 1E), representing one of the most significant enhancements among previously reported GPCR constructs (14, 21, 33, 37). This result suggests that M1R-G_11_CT maintains an exceptionally stable active state.

### M1R-G_11_CT maintained active-state conformational stability

We used M1R-miniGsq as a control to assess the conformational stability of M1R-G_11_CT. M1R-miniGsq, a widely used construct in structural analysis, integrates miniGsq protein—which features a truncated alpha helical domain (AHD) and an α5 helix substitution from Gq protein—at the C-terminus of the receptor. M1R-miniGsq exhibited a Kd value of 0.4 nM for [³H]-NMS (SI Appendix, Fig. S1G). Of note, M1R-miniGsq exhibits a biphasic binding curve (Khigh = 0.2 nM, Klow = 280 nM), indicating that only a fraction of receptor displays an increase in agonist affinity induced by G protein coupling. Similar biphasic curves were reported in other GPCR–G protein complexes (14, 38). In contract, M1R-G_11_CT exhibits a monophasic binding curve (Ki = 0.1 nM) (Fig. 1F), indicative of a homogeneous high-affinity agonist state akin to G-protein coupled M1R.

### The structure of M1R-G_11_CT represents agonist-free active state

As described above, M1R-G_11_CT demonstrated excellent biochemical properties throughout the purification process, even in the absence of agonist supplementation. We subsequently determined the cryo-EM structure of M1R-G_11_CT without ligand at the resolution of 3.62 Å (Fig. 2A and SI Appendix, Fig. S2). Two-dimensional class averages clearly delineated the secondary structural features within both the transmembrane domain and the designed fusion protein, confirming the proper folding of the engineered receptor (Fig. 2B). The density map of the orthosteric pocket lacks the complete ligand density previously observed in the active-state M1R bound to iperoxo, supporting that the receptor is in a ligand-free state. However, a residual, undefined density remains at the position of its tertiary amine group, likely contributed by a cation in the buffer, such as sodium (SI Appendix, Fig. S3A). Moreover, the orthosteric pocket exhibits a conformation similar to that of the active-state M1R bound to the agonist iperoxo (Fig. 2C). And the experimentally determined structure exhibited remarkable congruence with our computational prediction (SI Appendix, Fig. S3B). This structural correspondence between experimental and predicted models underscores the reliability of our protein engineering approach.

**Fig. 2.**
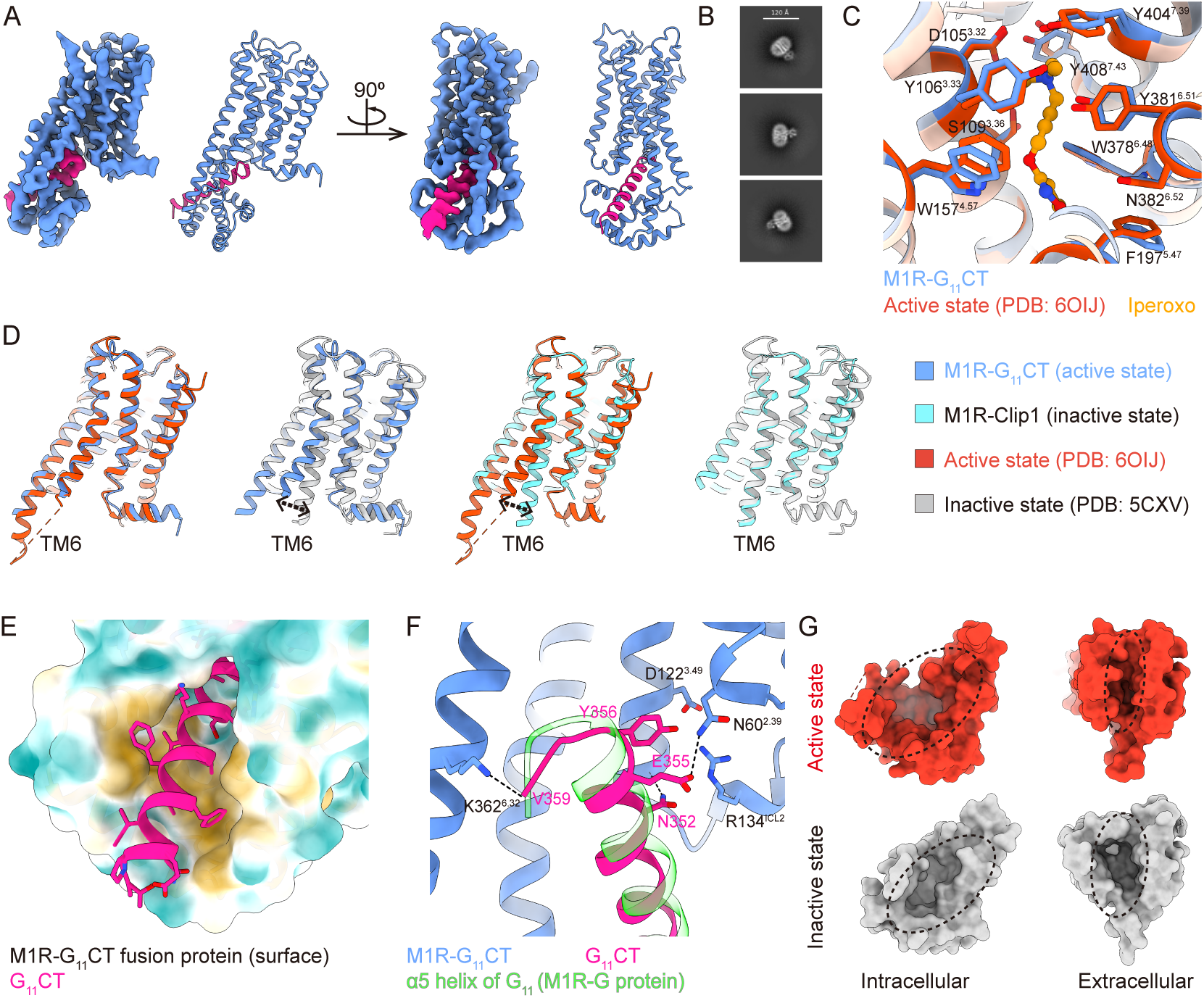
The structure of M1R-G_11_CT reveals its agonist-free active state. (A) Overall cryo-EM density map and atomic model of M1R-G_11_CT. (B) Representative 2D class averages of M1R-G_11_CT particles, highlighting the transmembrane helices of receptor and the designed fusion protein. (C) The orthosteric binding pocket in M1R-G_11_CT closely resembles that of the previously resolved active-state structure bound to iperoxo (PDB: 6OIJ), confirming its stabilization in an active receptor conformation. (D) Structural alignment of M1R-G_11_CT and M1R-Clip1 with both active-state M1R (PDB: 6OIJ) and inactive-state M1R (PDB: 5CXV) structures, emphasizing the displacement of TM6. (E) The de novo designed fusion protein forms a hydrophobic binding pocket that accommodates the N-terminus of G_11_CT. (F) The C-terminus of G_11_CT adopts a binding pose analogous to the α5 helix of the G protein in the previously resolved M1R-G protein complex bound to iperoxo (PDB: 6OIJ), further validating its functional mimic of G protein interactions. (G) Top and bottom views comparing the extracellular and intracellular pockets of active-state and inactive-state M1R.

Structure comparison was conducted between two engineered receptors (the inactive-state M1R-Clip1 and the active-state M1R-G_11_CT) and their respective state-specific structures to evaluate the design outcomes (Fig. 2D). For M1R-Clip1, TM6 exhibits an inward displacement characteristic of the inactive state (PDB: 5CXV), differing from the outward movement observed in the active state (PDB: 6OIJ). Conversely, TM6 in M1R-G_11_CT retains an outward displacement, maintaining the angle and position analogous to the active state, distinct from the inward displacement of the inactive state. The designed fusion protein of M1R-G_11_CT forms a well-defined hydrophobic pocket that accommodates the G_11_CT (Fig. 2E). The position and orientation of G_11_CT within the intracellular pocket of M1R closely mirror those observed in the M1R-G_11_ protein complex (Fig. 2F). Of note, the G_11_CT of ligand-free M1R-G_11_CT can still precisely inserts into the intracellular pocket of the receptor, indicating that the active state of M1R-G_11_CT is preserved independently of agonist activation. Taken together, we have successfully validated engineered M1R proteins with conformational stability in both the inactive state (M1R-Clip1) and the active state (M1R-G_11_CT) through biochemical and structural analyses, enabling binding-based drug screening. Importantly, M1R-G_11_CT maintains active state even under agonist-free condition, representing an advancement compared to previous strategies for generating active-state receptor.

### Intracellular orthogonal screening for extracellular nanobodies

In the realm of binding-based drug screening, extracellular antibody screening targeting Class A GPCRs has long been a challenge. The difficulty stems from the inherent asymmetry in the solvent-accessible surface area (SASA) between the intracellular and extracellular regions of class A GPCRs. The M1R receptor serves as an example, where the intracellular SASA in its inactive state exceeds that of the extracellular region. Furthermore, upon receptor activation, the intracellular SASA expands while the extracellular SASA becomes more contracted (Figure. 2G). This results in a preferential enrichment of antibodies targeting the intracellular region, whereas antibodies specific to the extracellular region—especially those recognizing the active state—are often disadvantaged and routinely excluded during conventional binding-based antibody screening.

To address these challenges, we define intracellularly orthogonal receptors as those sharing the identical extracellular structures but possessing distinct intracellular regions. Examples of such receptor pairs include the inactivated atropine-bound M1R-ICL3 and M1R-Clip1, as well as the activated pair iperoxo-bound M1R-G_11_CT and M1R-G_11_CTb. By employing these intracellularly orthogonal receptors in iterative rounds of cross-screening, it is feasible to directionally enrich antibodies targeting the extracellular pocket. In this study, we screened for nanobodies specific to the inactive and active states using the aforementioned receptor pairs as antigens, respectively, from a synthetic nanobody library displayed on yeast surface (Fig. 3 A, B and C). Several rounds of Magnetic-Activated Cell Sorting (MACS) and Fluorescence-Activated Cell Sorting (FACS) were performed with progressively reduced receptor concentration to enhance the affinity of the selected nanobodies (SI Appendix, Fig. S4). Although we are able to generate stable active or inactive state M1R in the apo state using fusion proteins, the specific tyrosine lid structure within the receptor (21) hinders the screening of orthosteric nanobodies targeting M1R. As a result, our project focuses on identifying allosteric nanobodies targeting the receptor.

**Fig. 3.**
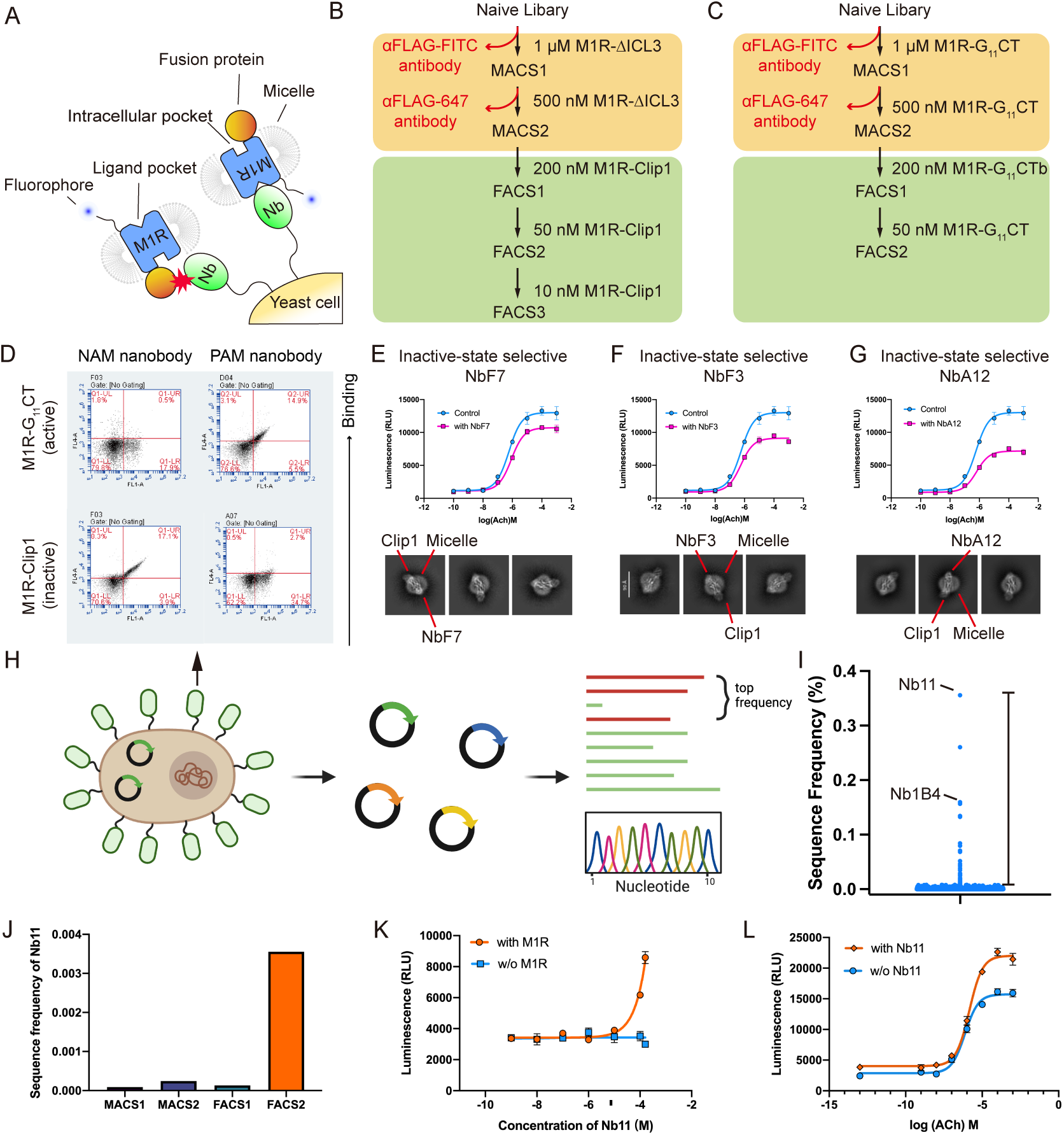
Intracellular orthogonal screening for extracellular nanobodies (A) Engineered proteins M1R-G_11_CT, M1R-G_11_CTb and M1R-Clip1 demonstrate dual functionality: stabilizing distinct receptor conformations while preferentially enriching extracellular pocket-targeting nanobodies over intracellular-targeting counterparts during yeast display screening. (B) Schematic representation of the intracellular orthogonal screening approach for isolating inactive-state selective nanobodies. (C) Schematic representation of the intracellular orthogonal screening approach for isolating active-state selective nanobodies. (D) Yeast surface staining analysis, performed with 10 nM fluorescence-labeled receptors, validates the conformational selectivity of nanobodies, demonstrating their capacity to differentiate between receptor states. (E-G) Cell-based functional assays and 2D class averages of inactive-state selective nanobodies (NbF7, NbF3, and NbA12) reveal that these extracellular binding nanobodies attenuate the endogenous agonist signaling efficacy (Emax) at 100 μM. (H) Schematic representation of the next-generation sequencing (NGS) workflow employed for nanobody identification. (I) Nb11 and Nb1B4 demonstrated predominant enrichment profiles during active-state selective screening, emerging as the most abundant nanobody variants. (J) The progressive enrichment of Nb11 throughout the screening process. Functional characterization demonstrates that Nb11 exhibits partial agonist efficacy (K) and potentiates the maximal response of endogenous agonist signaling at 100 μM (L) in Glo-sensor assays. Data represent mean values ± SEM derived from three independent experimental replicates.

Following screening, the single colonies of yeast were stained using fluorescently labeled atropine-bound M1R-Clip1 and iperoxo-bound M1R-G_11_CT proteins to identify conformation-specific nanobodies (Fig. 3D), and the inactive-state selective nanobodies (NbF3, NbF7, NbA12) and the active-state selective nanobody (Nb1B4) were selected for further validation. Following the purification of nanobodies, a Glo-sensor-based signaling assay was performed to assess their pharmacological properties. Among the nanobodies selective for the inactive state, NbF7, NbF3, and NbA12 were identified to reduce the Emax of acetylcholine (ACh), the endogenous agonist. Furthermore, in the β-arrestin recruitment assay, we demonstrated that NbA12 exhibits a certain degree of selectivity among the five subtypes (SI Appendix, Fig. S5A). Additionally, these nanobody features are localized on the side opposite to Clip1 in the two-dimensional classification, indicating that these nanobodies bind extracellularly (Fig. 3 E, F and G). However, we could not detect the agonist activity of Nb1B4, likely because the protein tends to aggregate after purification, which impedes the assessment of its function. To identify additional active-state selective nanobodies, we performed next generation sequencing (NGS) to analyze the nanobodies’ enrichment pattern of the active-state screening group (Fig. 3H). The most enriched clone, Nb11, was progressively accumulated throughout the screening process (Fig. 3 I and J). Pharmacological assays demonstrated that Nb11 directly activates the M1R, albeit with low potency and efficacy compared to iperoxo, while moderately enhances the Emax of Ach (Fig. 3 K and L and SI Appendix, Fig. S5B).

### Structure insight of nanobody-M1R complexes

To better understand the molecular binding mode of the conformationally selective nanobodies, we attempted to resolve the complex structures of these nanobodies with the receptor. Unfortunately, the strong orientation preference of NbF3-bound and NbF7-bound complex samples prevented us from obtaining their three-dimensional structures. Finally, we successfully determined the cryo-EM structures of the active-state selective Nb1B4 bound to M1R-G_11_CT complex with iperoxo, as well as the inactive-state selective NbA12 bound to M1R-Clip with atropine at global resolutions of 2.88 Å and 3.29 Å, respectively (Fig. 4 A and B, and SI Appendix, Fig. S6 and S7). For Nb1B4, the structure revealed unambiguous electron density for Nb1B4-M1R interface. However, the NbA12-M1R samples still suffers from preferred orientation and the nanobody part exhibits discontinuous electron density, which prevented us from building a complete nanobody model. Despite this limitation, we were able to approximately localize the position of its CDR3 based on the available density (SI Appendix, Fig. S8A). The overall maps reveal that Nb1B4 and NbA12 share a similar scaffold binding pose, albeit with a rotational difference, when compared to MT7, a negative allosteric modulator (NAM) for M1R (11) (Fig. 4 C and D). This observation suggests that the extracellular vestibule of the muscarinic receptor is sufficiently spacious to accommodate diverse binding orientations, thereby facilitating the development of novel modulators targeting the receptor. Muscarinic receptors have long served as prototypic model systems for elucidating allosteric modulation of GPCR signaling. These structures confirmed that the CDR3 loops of Nb1B4 and NbA12 both occupy the conventional allosteric vestibule of M1R, located above the orthosteric site (Fig. 4 E and F). And the binding poses of the orthosteric ligands are similar to those observed in previously reported structures without nanobodies (SI Appendix, Fig. S8 B and C).

**Fig. 4.**
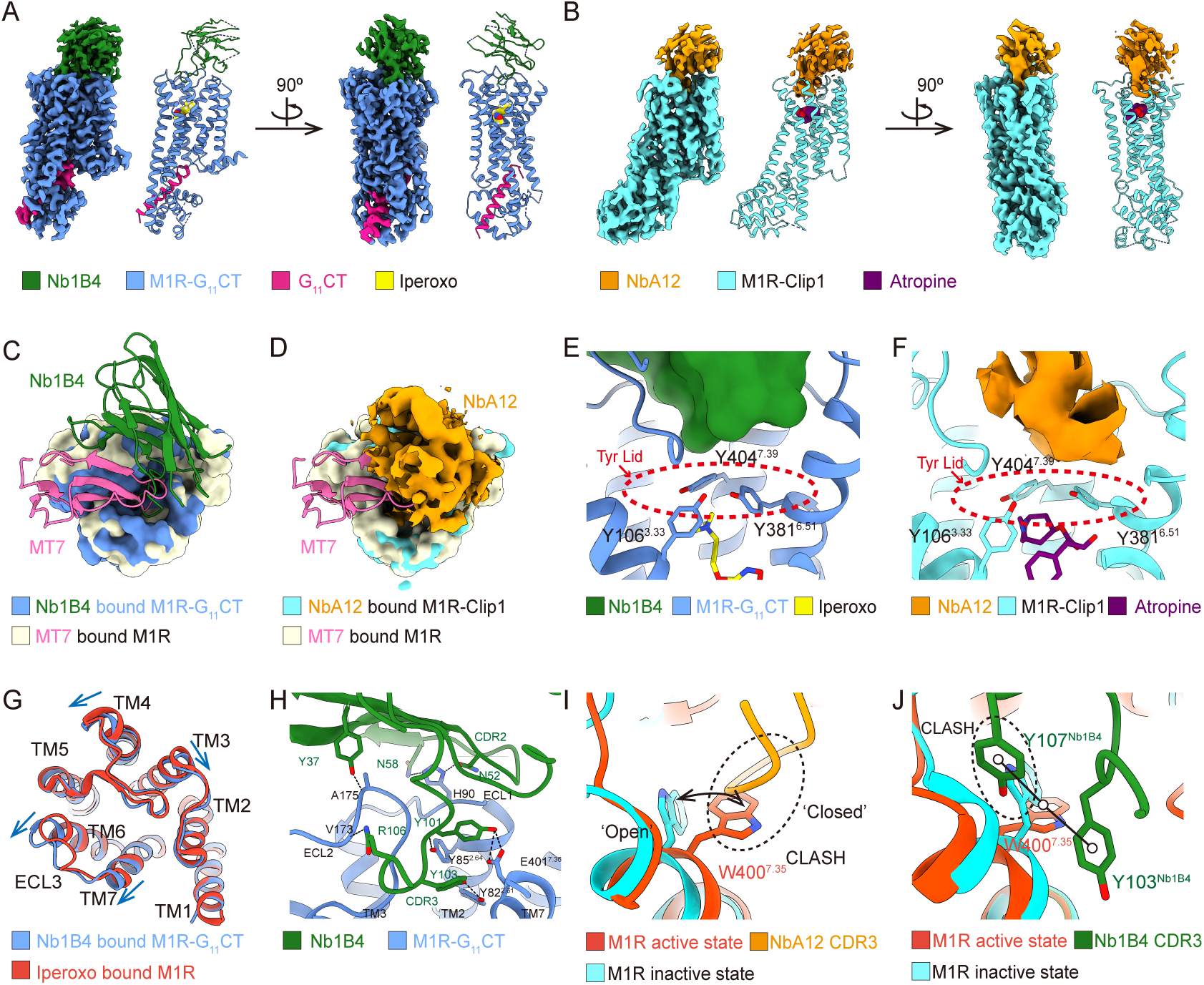
Structure insight of nanobody-M1R complexes (A) Overall cryo-EM density map and atomic model of the Nb1B4 and M1R-G_11_CT complex bound to iperoxo. (B) Overall cryo-EM density map and refined atomic model of the NbA12 and M1R-Clip1 complex bound to atropine. The binding orientation of (C) Nb1B4 and (D) NbA12 are similar but distinct from that of MT7 (MT7 bound M1R, PDB: 6WJC). (E) Nb1B4 and (F) NbA12 bind to the allosteric pocket of M1R, positioned above the orthosteric pocket and tyrosine lid. (G) Nb1B4 binding induces outward displacement of TM6, TM7, and ECL3 compared to the iperoxo-bound M1R structure (PDB: 6OIJ). (H) Nb1B4 engages the extracellular vestibule of M1R through a broad network of polar interactions. (I) NbA12’s CDR3 region exhibits conformational selectivity by accommodating the open state of W400^7.35^ while sterically clashing with its closed state conformation. (J) Nb1B4’s CDR3 region, particularly residues Y107 and Y103, demonstrates conformational selectivity by stabilizing the closed state of W400^7.35^ while exhibiting steric clashes with its open state conformation.

Nb1B4 is the first nanobody identified to bind to the extracellular allosteric pocket of the active-state M1R with conformational selectivity (Fig. 3D). The high-resolution complex structure provides precise interaction details. Structure comparison between the M1R-Nb1B4-iperoxo complex and the nanobody-free M1R-iperoxo structure (PDB:6OIJ) reveals that Nb1B4 CDR3 insertion drives a slight outward movement of extracellular ends of TM6, TM7, and ECL3 (Fig. 4G). The Nb1B4 scaffold, along with its CDR2 and CDR3, engages in extensive polar interactions with the receptor’s extracellular surface. The intricate hydrogen bond network includes interactions between Y37^Nb1B4^ and R106^Nb1B4^ with the backbone of the receptor’s ECL2, hydrogen bonds between N58^Nb1B4^ and N52^Nb1B4^ with H90^ECL1^, and additional hydrogen bonds formed by Y101^Nb1B4^ with E401^7.38^ and Y85^2.64^, as well as Y103^Nb1B4^ with Y82^2.61^. This region has been identified in prior studies as a critical site for the function of allosteric modulators (11, 21, 39) (Fig. 4H).

### Conformational selectivity mechanism of nanobodies

Structural comparisons have revealed several key factors influencing conformational selectivity. Overall, NbA12 binds to a broad pocket, similar to the allosteric pocket of the inactive state, whereas Nb1B4 binds to a narrow pocket, resembling the allosteric pocket of the active state (SI Appendix, Fig. S8D). Specifically, the CDR3 region of NbA12 is positioned closer to TM5, while that of Nb1B4 is located nearer to TM2 (SI Appendix, Fig. S8E). This spatial difference in CDR3 positioning underlies their distinct conformational preferences. The conformational changes in key amino acid residues modulate the morphology of the receptor’s allosteric pocket between different states, with W^7.35^ playing a critical role. This residue undergoes a flip between the two states, a phenomenon highly conserved across the acetylcholine receptor family (SI Appendix, Fig. S8F). In the active state, W^7.35^ adopts a “closed” conformation, while in the inactive state, it assumes an “open” conformation (Fig. 4I).

For NbA12, the “closed” conformation of W400^7.35^ clashes with its backbone, restricting its binding to the inactive state receptor (Fig. 4I). In contrast, for Nb1B4, the interaction interface is closer to TM2, providing sufficient space around W400^7.35^ to accommodate its “closed” conformation. Additionally, Y107^Nb1B4^ clashes with W^7.35^ in the “open” state, while Y103^Nb1B4^ forms hydrophobic interactions with W400^7.35^ in the “closed” state, stabilizing the latter conformation (Fig. 4J and SI Appendix, Fig. S8G). These structural and interactional nuances collectively enable Nb1B4 to selectively bind to the active state receptor. Notably, the CDR3 of Nb1B4 forms a robust and conserved hydrogen bond network with TM7 and TM2, a feature also observed in the structures of positive allosteric modulators (PAMs) targeting M2R and M4R, such as LY2119620, LY2033298, and VU0467154 (SI Appendix, Fig. S8H) (21). This conserved interaction underscores the critical role of this site as a key allosteric binding hotspot within the muscarinic receptor family.

## Discussion

G protein-coupled receptors (GPCRs) have long been recognized as crucial therapeutic targets, with both pharmaceutical industry and academic researchers dedicating decades of effort to novel ligand discovery. In recent years, binding-based drug discovery has emerged as a transformative approach in GPCR-targeted therapeutic development, demonstrating remarkable success through multiple breakthrough achievements. This approach has enabled significant milestones, including the application of DEL technology in identifying the first negative allosteric modulator and the first positive allosteric modulator for the β2AR, and the utilization of hybridoma screening in the development of Erenumab, which stands as the first FDA-approved GPCR-targeting antibody drug (40–43). The exceptional potential of binding-based drug discovery lies in its inherent advantages: the capability to target both orthosteric and allosteric binding pockets, the capacity to explore an extensive chemical space, and the facilitation of diverse ligand discovery encompassing antibodies, peptides, and small molecules.

It is widely recognized that obtaining a stable, purified receptor—exhibiting both thermal and structural stability—is a crucial prerequisite for successful binding-based screening. In this study, we investigated the feasibility of leveraging AI-driven approaches to design tool proteins that facilitate binding-based screening efforts. Building on our two previous studies, we refined the protein design method and successfully engineered M1R-Clip1, stabilized in the inactive state, and M1R-G_11_CT, stabilized in the active state (32, 33). Notably, both the ligand-free tool proteins (M1R-Clip1 and M1R-G_11_CT) exhibit significantly enhanced biochemical stability, a crucial feature for effective binding-based screening applications. Among these, M1R-G_11_CT stands out with exceptional performance, demonstrating an approximately 600-fold improvement in agonist affinity. Remarkably, the agonist-independent active state of the engineered protein preserves the intact extracellular vestibule of the receptor, particularly the unoccupied orthosteric pocket for binding-based screening. Compared to our previous approach which relied on T4 lysozyme (T4L) fusion and disulfide bond stabilization (33), the novel workflow avoids the need of extensive construct screening. More significantly, it enables the efficient generation of a series of fusion proteins sharing the same scaffold but featuring distinct sequences, which we term intracellular orthogonal receptors. This advancement is pivotal for reducing screening noise attributable to the fusion protein itself, thereby enhancing the precision and reliability of the screening process.

Based on the designed proteins, we performed a proof-of-concept nanobody screening from a synthetic yeast surface display library. Antibodies targeting GPCRs have attracted considerable attention due to their exceptional selectivity and favorable pharmacokinetic properties. To overcome the challenges associated with conformationally selective antibody screening against the extracellular domains of GPCRs, several innovative approaches have been proposed. The core concept involves introducing or blocking diverse surfaces of the target protein to enrich nanobodies that specifically recognize particular epitopes (9, 44, 45). Our intracellular orthogonal screening strategy aligns with this rationale but offers significantly enhanced versatility and scalability. By utilizing tool proteins with distinct intracellular sequences but identical extracellular regions in each screening round, we successfully enriched nanobodies specifically targeting the extracellular domain. This approach ultimately enabled us to isolate a series of conformationally selective nanobodies against M1R.

The muscarinic receptor, a central therapeutic target for neuropsychiatric disorders like Alzheimer’s and schizophrenia under the cholinergic hypothesis, has driven decades of research into selective agonists and PAMs. In this study, we introduce the first series of extracellular nanobodies targeting M1R. Cryo-EM analysis reveals that these nanobodies bind to an allosteric pocket that undergoes conformational changes during receptor activation—a site shared by LY2119620 and muscarinic toxin MT7. Stabilizing this pocket to maintain the receptor conformation may represent a general mechanism for the allosteric modulation of muscarinic receptors. Moreover, these nanobodies are readily amenable to affinity maturation or biochemical optimization, enabling the enhancement of their binding affinity and functional activity in future research.

## Materials and Methods

### Computational design of M1R_G_11_CT and M1R_G_11_CTb

M1R-G_11_CT was engineered through an integrative computational design approach including RFdiffusion, ProteinMPNN and AlphaFold2 (46–48), following a well-established protocol previously employed for click fusion protein (Clip) design (32). The design process commenced with the extraction of the active-state M1R and α5 helix of G_11_ protein complex from M1R-G_11_ protein complex (PDB: 6OIJ) as a substructure template. Subsequently, the ICL3 region of M1R was removed, and two residues (T215^5.65^from TM5 and A364^6.34^ from TM6) were selected as fusion site for backbone generation of the fusion protein, while maintaining the α5 helix of G_11_ protein as an independent chain. The backbone architecture of the fusion protein was initially generated using RFdiffusion, followed by the side chain design through ProteinMPNN. Structural refinement was performed through an iterative process, wherein regions exhibiting low pLDDT scores predicted by AlphaFold2 were locally refined while preserving high-confidence backbone conformations with superior pLDDT scores. After multiple cycles of refinement, the amino acid sequence demonstrating high pLDDT scores was selected, and the corresponding DNA sequence encoding the fusion protein was synthesized for experimental characterization. Two protein constructs, M1R-G_11_CT, M1R-G_11_CTb, which share an identical backbone but have distinct side chains, were generated for in vitro screening of nanobodies.

### Expression and purification of the receptors

The M1R constructs (M1R, M1R-ΔICL3, M1R-G_11_CT, M1R-G_11_CTb, and M1R-Clip1) were cloned into pFastBac vector for expression via the Bac-to-Bac Baculovirus system (Invitrogen). Sf9 cells were infected at a density of approximately 4.0 × 10⁶ cells/mL and harvested 48 hours post-infection. M1R-ΔICL3 expression was enhanced by supplementing 10 μM atropine during infection. Receptor purification was performed using Ni-NTA, Flag affinity chromatography, and size-exclusion chromatography (SEC) as described previously (6). Briefly, cell pellets were lysed in lysis buffer (10 mM HEPES, pH 7.5, 1 mM EDTA, 160 μg/mL benzamidine, 100 μg/mL leupeptin) for 30 minutes at room temperature. Following complete lysis, cell membranes were collected by centrifugation and solubilized using solubilization buffer (20 mM HEPES, pH 7.5, 750 mM NaCl, 1% dodecyl maltoside (DDM), 0.2% Na Cholate, 0.03% cholesterol hemisuccinate (CHS), 30% glycerol, 5 mM imidazole, 2 mM MgCl2, 160 μg/mL benzamidine, 100 μg/mL leupeptin, 2 mg/mL iodoacetamide and benzonase), with continuous stirring at 4°C for 2 hours. After high-speed centrifugation, the supernatant was incubated with Ni-NTA resin for 2 hours at 4 °C. The Ni-NTA resin, bound with receptor, was washed and then eluted with elution buffer (20 mM HEPES pH 7.5, 750 mM NaCl, 0.1% DDM, 0.02% Na Cholate, 0.03% CHS, 200mM imidazole). Next, the elution from Ni-NTA resin was incubated with M1-Flag affinity resin with addition of 2 mM CaCl2 and the detergent was gradually exchanged from DDM to 0.01% lauryl maltose neopentyl glycol (MNG). The receptor was eluted with 0.2 mg/mL Flag peptide and 5 mM EDTA, followed by further purification on a Sephadex 200 Increase 10/300 GL column (Cytiva) with a running buffer (20 mM HEPES pH 7.5, 0.01% MNG, 0.001% CHS, 100 mM NaCl). Finally, the purified receptor was concentrated using 50 kDa molecular weight cutoff Amicon centrifugal filters (Millipore).

### Cryo-EM grid preparation

Purified M1R-G_11_CT (4 μL) was deposited onto glow-discharged nickel-titanium grids (Ni-Ti R1.2/1.3, 300 mesh), while purified M1R-nanobody complexes (4 μL) were deposited onto glow-discharged Au Quantifoil grids (Au R1.2/1.3, 200 mesh). The grids were then gently blotted with Whatman No. 1 qualitative filter paper in a Vitrobot Mark IV (Thermo Fisher) chamber, which was maintained at 8 °C and 100% humidity. Blotting parameters were set to a blot time of 4 seconds and a blot force of four. Immediately after blotting, the grids were rapidly vitrified by plunging into liquid ethane.

### Data collection and processing

Cryo-EM data of apo-state M1R-G_11_CT (1960 micrographs), M1R-G_11_CT-iperoxo-Nb1B4 complex (1950 micrographs), M1R-Clip1-atropine-NbA12 complex (3668 micrographs) were collected using Titan Krios G3i TEM (Thermo Fisher) equipped with an automated data collection (AutoEMation) (49). Data processing was conducted using a standardized workflow in cryoSPARC (v4.5.1) (50). Initial 2D classifications were carried out, and the highest-quality 2D averages were selected for 3D reconstruction. These averages were subsequently used as templates for particle picking across the entire dataset. Following multiple rounds of heterogeneous refinement and non-uniform refinement, a 3D volume with clear secondary structures was obtained. To focus on the transmembrane domain, detergent micelles in the 3D volume were masked out, enabling focused classification. After several iterations of heterogeneous refinement and 3D classification, high-resolution volumes were ultimately achieved. The final structure reconstructions utilized 155,176 particles for apo-state M1R-G_11_CT, 191,008 particles for the M1R-G_11_CT-iperoxo-Nb1B4 complex, and 254,687 particles for the M1R-Clip1-atropine-NbA12 complex, yielding resolutions of 3.62 Å, 2.88 Å, and 3.29 Å, respectively.

### Model building and refinement

The initial structural models of M1R-G_11_CT, M1R-Clip1, Nb1B4 and NbA12 were generated from AlphaFold2 predictions. Coordinates and chemical constraints for atropine and iperoxo were created using Phenix.elbow (1.20.1-4487) (51). These models were then fitted into the Cryo-EM density maps using UCSF ChimeraX-1.3 (52), followed by manual adjustments and refinements in COOT-0.9.8.7 (53). Final structural refinement and validation were carried out using PHENIX (54).

### Cell surface staining

Cell surface staining was performed to assess receptor expression levels. Briefly, transfected cells were resuspended in phosphate buffer solution (PBS) containing 2 mM CaCl2 and incubated with Alexa-488 conjugated anti-Flag antibody (diluted in HBSS at a ratio of 1:300, Thermo Fisher, Cat # MA1-142-A488) for 15 minutes at room temperature in the dark. Then the cells were washed twice with PBS containing 2 mM CaCl2. Non-specific binding was evaluated using negative control samples containing 0.2 mg/mL Flag peptide. Fluorescence measurements were then performed using a BD Accuri C6 flow cytometer (BD Biosciences), with excitation at 488 nm and emission at 519 nm. Cells were gated by cell size and granularity to ensure accurate analysis.

### Radioligand binding assays

The M1R constructs, including M1R-G_11_CT, M1R-Clip1, M1R-ΔICL3 and M1R-miniGsq, were expressed in either Sf9 insect cells or HEK293F mammalian cells. Membrane fractions were prepared by harvesting 50 mL of cell culture, followed by homogenization in 8 mL of ice-cold lysis buffer (20 mM Tris, pH 7.5, 1 mM EDTA). The homogenate was subjected to sequential centrifugation: first at 800 ×g for 10 min to remove cellular debris, then the supernatant was centrifuged at 18,000 ×g for 20 min to pellet membranes. The final membrane fraction was resuspended in binding buffer (20 mM HEPES, pH 7.5, 100 mM NaCl). For saturation binding assays, membranes were incubated with [^3^H]-NMS (0-4 nM) in binding buffer (20 mM HEPES, pH 7.5, 100 mM NaCl, 5 mM MgCl2, and 0.1% BSA) for 1 hour at room temperature. Atropine (10 μM) was used to define nonspecific binding. For competition binding experiments, membranes were incubated with a constant concentration of [³H]-NMS alongside increasing concentrations of atropine or iperoxo for 60 minutes at room temperature to determine ligand binding affinities for M1R variants. Binding reactions were rapidly filtered through GF/B filters using Brandel 48-well harvester. Filters were washed three times with ice-cold binding buffer and immersed in scintillation fluid. Bound radioactivity was quantified using a liquid scintillation counter (MicroBeta2, PerkinElmer), and the data were analyzed with Prism 9 (GraphPad).

### Discovery of the conformationally selective nanobodies targeting M1R

The synthetic nanobody library displayed on the yeast surface was generously provided by Drs. A. C. Kruse (Harvard University) and A. Manglik (University of California San Francisco) (9). Yeast cells were cultured in tryptophan dropout (-Trp) medium supplemented with 2% (w/w) glucose at 30 °C. Nanobody expression was induced by transferring the cells to - Trp medium containing 2% (w/w) galactose, followed by culturing for 2 days at 25°C. Nanobody expression levels were assessed by staining with Alexa Fluor-488-conjugated anti-HA antibody (Cell Signaling Technology) and analyzed via flow cytometry using an Accuri C6 (BD Biosciences).

Nanobody clones targeting the different purified FLAG-tagged M1R constructs were enriched through two rounds of magnetic-activated cell sorting (MACS) and two or three rounds of fluorescence-activated cell sorting (FACS) using FACSAria II (BD Biosciences) in selection buffer (20 mM HEPES pH 7.5, 100 mM NaCl, 0.05% MNG, 0.005% CHS, 2 mM CaCl2, 0.1% (w/v) bovine serum albumin and 0.2% maltose), as previously reported (See Fig.3 B and C) (55). For the first and second rounds of MACS, 5×10⁹ and 2×10⁸ induced yeast cells were used as input, respectively. The yeast cells were washed and precleared by incubating with Alexa Fluor-FITC-conjugated anti-FLAG antibody and anti-Alexa Fluor-FITC microbeads (Miltenyi) (or Alexa Fluor-647-conjugated anti-FLAG antibody and anti-Alexa Fluor-647 microbeads), followed by passage through an LD column (Miltenyi) to remove yeast displaying nonspecific nanobodies. The flow-through yeast cells were washed with selection buffer, and then incubated with FLAG-tagged M1R construct bound to ligand (atropine for inactive-state selective nanobody screening, iperoxo for active-state selective nanobody screening), Alexa Fluor-FITC-conjugated anti-FLAG antibody and anti-Alexa Fluor-FITC microbeads (or Alexa Fluor-647-conjugated anti-FLAG antibody and anti-Alexa Fluor-647 microbeads). After incubation at 4°C for 30 minutes, the yeast cells were loaded onto an LS column, washed with selection buffer and eluted by plunger.

Subsequently, several rounds of FACS were performed using FACSAria II (BD Biosciences) (See Supplementary Fig.4). For each round, 2×10^7^ yeast cells were stained with Alexa Fluor-488-conjugated anti-HA antibody (Cell Signaling Tech), Alexa Fluor-647-conjugaed anti-FLAG antibody and FLAG-tagged M1R construct bound with ligand (atropine for inactive-state selective nanobody screening, iperoxo for active-state selective nanobody screening) in the selection buffer.

After screening, the sorted yeast cells were diluted and plated on -Trp agar plates, and single colonies were picked and cultured as clonal populations in deep 96-well plates for nanobody binder screening. For yeast surface staining, the nanobody displayed yeast cells were stained with the Alexa Fluor-488-conjugated anti-HA antibody (Cell Signaling Tech), purified FLAG-tagged M1R and Alexa Fluor-647-conjugated anti-FLAG antibody in the selection buffer, and analyzed by Accuri C6 (BD Biosciences). The double-positive staining yeast cells among the anti-HA positive population were selected for sequencing analysis.

### Expression and purification of nanobodies

Selected yeast clones were sequenced and the corresponding nanobody sequences were subcloned into the pET26b periplasmic expression vector featuring an N-terminal pelB signal peptode and a C-terminal histidine tag, and then transformed into BL21(DE3) *Escherichia coli*. Cells were grown in Terrific Broth (TB) medium supplemented with 2 mM MgCl2, 0.1% glucose, and 50 µg/ml kanamycin at 37°C until reaching an OD600 of 1.0. Protein production was induced with 1 mM IPTG, and the culture was incubated at 20°C for 20 hours with shaking. Periplasmic extraction was performed via osmotic shock, cells were resuspended in cold hypertonic buffer (0.2 M Tris-HCl, pH 8.0, 0.5 mM EDTA, 0.5 M sucrose) for 1 hour at 4°C, then diluted 4-fold with ice-cold water for additional hypotonic treatment. After high-speed centrifugation, the supernatant was purified using Ni-NTA resin followed by SEC with running buffer (20 mM HEPES, pH7.5, 100 mM NaCl). In addition to the *E.coli* expression system, some nanobodies such as NbF3, NbF7, NbA12 and Nb1B4 were expressed in HEK293F transient expression system. These nanobodies were subcloned into pcDNA3.4 vector with an N-terminal Mouse Ig Heavy chain signal peptide and a C-terminal histidine tag, cells were transfected with Polyethylenimine (PEI) at a density of 2.0 × 10^6^ cells/mL and cultured at 37 °C for 3–4 days. The culture supernatants were harvested for nanobody purification using Ni-NTA resin and SEC. Finally, the purified nanobodies were concentrated to 10-40 mg/mL using 10 kDa molecular weight cutoff Amicon centrifugal filters (Millipore).

### Next generation sequencing (NGS)

To track sequence evolution throughout the selection process, nanobody-encoding regions were PCR-amplified from yeast plasmid populations using primers (seq-F: 5’-CCTGCGCCTGAGCTGCG-3’, seq-R: 5’-GCTGCTCACGGTCACCTGGG-3’). The PCR products were sent to Sangon Biotech (Shanghai, China) Co.Ltd for sequencing library preparation and subsequent paired-end sequencing (PE300) on the Illumina platform, with raw reads processed through NGmerge for pair merging. Quality-filtered sequences were translated and analyzed for amino acid frequency distributions using custom scripts.

### Glo-sensor signaling assay

A Gsq chimera, constructed by replacing the last 15 amino acids of C-terminus of Gs protein with the last 15 amino acids of Gq protein, was used to measure Gq signaling using Glo-sensor assay (56). To perform the assay, HEK293T cells were seeded in a six-well plate and allowed to grow to 50–80% confluence prior to transfection. A plasmid mixture containing 250 ng M1R-wt, 250 ng Gsq, and 3750 ng pGloSensor^TM^-22F plasmid (Promega) in 250 µl of Opti-MEM (Gibco) was transfected into cells using 12.5 uL 1 mg/mL PEI. 24 hours post-transfection, cells were harvested from the plate and resuspended in 5 mL Hanks’ Balanced Salt Solution (HBSS) supplemented with 20 mM HEPES, pH 7.5 and 150 µg/mL luciferin, and then transferred to a 96-well plate at 70 µL per well. After incubation at 37°C for 1 hour followed by an additional hour at room temperature, 20 µL of different concentrations of nanobodies and 10 µL of acetylcholine (Ach) were added, and luminescence counts were measured by Ensight^TM^ plate reader (PerkinElmer). For the inactive-state selective nanobodies, a 20-minute pre-incubation with nanobodies was performed before Ach addition. Data analysis was conducted using Prism 9 (GraphPad).

### NanoBiT based β-arrestin recruitment assay

NanoBiT based β-arrestin recruitment assays were performed following a previously published protocol with minor modifications (57). Briefly, Hek293T cells were transfected with corresponding plasmids using PEI as transfection reagent at a DNA:PEI ratio of 1:3. 1.3 μg of receptor-SmBiT (with M2R-M5R constructs containing the V2Rpp sequence at the C-terminus, except for the M1R-wt construct) and 1.3 μg of LgBiT-β-arrestin1 were used. After 18-20 hours of transfection, cells were harvested in DMEM, centrifuged and resuspended in 5 mL HBSS buffer supplemented with 10 mM HEPES, pH7.5 and coelenterazine. The cells were then seeded in a white, flat-bottom, 96-well plate at 70 μL per well. The plate was incubated at room temperature for 60 minutes to allow for equilibration. Nanobodies were added at the final concentration of 100 μM and incubate for 20 minutes, followed by recording the luminescence counts as the basal reading using Ensight^TM^ plate reader (PerkinElmer). The cells were subsequently simulated with varying doses of Ach (10 µL per well) prepared in buffer (20 mM HEPES, pH7.5, 100 mM NaCl), and luminescence was recorded. Data analysis was conducted using Prism 9 (GraphPad).

## Acknowledgments

This work was supported by Beijing Frontier Research Center for Biological Structure, and Tsinghua-Peking Center for Life Sciences, Tsinghua University (X.L.), by National Natural Science Foundation of China (Grant 32122041 to X.L.) and by Tsinghua University Initiative Scientific Research Program (X.L.). We gratefully acknowledge Drs. A. C. Kruse (Harvard University) and A. Manglik (University of California San Francisco) for supplying the synthetic nanobody library.

## Author Contribution

X.Z. performed the protein design, construction with the help of K.G. X.Z., N.J. and K.G. and H.M. performed protein expression and purification. K.G. prepared the Cryo-EM samples. X.Z. and K.G. collected the Cryo-EM data. X.Z. and K.G. performed structure determination and refinement. X.Z. and X.L. performed model building. K.G. and X.Z. performed the radioligand binding assay. X.Z. and J.N. performed the nanobody screening, and X.Z. and K.G. performed the characterization of nanobodies. X.L. supervised overall project. X.Z., K.G. and X.L. wrote the manuscript. All authors discussed the results and commented on the manuscript.

## Notes

### Competing Interest Statement

The authors have declared no competing interest.

